# Lifespan in Bats: Enigmatic Longevity and Evolutionary Stasis

**DOI:** 10.1101/2025.07.29.667414

**Authors:** Anastasiia Lukianchuk, Igor Dzeverin

## Abstract

Bats are known to live significantly longer than other mammals of similar body size – a fact that continue to puzzle researchers, as the underlying reasons behind bats’ extended lifespan remain poorly understood. Our study aims to identify key morphometric and life-history traits that contribute to extreme longevity of bats and to describe the evolutionary patterns of bat lifespan. Using phylogenetically informed regression with dummy variables and dataset of 108 bat species, we examined the effects of body length, body weight, forearm length, diet, reproduction rate, hibernation, and climatic preference on longevity. Our analysis revealed that lifespan in bats is influenced by their body size (the proxy for which is the forearm length), reproduction rate, some specific diets (hematophagy and omnivory), and climate, while other traits showed no noteworthy effects. Moreover, we fitted several likelihood models for the evolution of forearm length, body length, body mass and lifespan. All these traits best followed an Ornstein–Uhlenbeck model. As the result, lifespan appeared to be a highly constrained trait with an estimated evolutionary optimum of approximately 13.5 years across Chiroptera. These findings support the idea that longevity may have been present in ancestral bat lineages and is shaped by the combination of ecological and morphological factors.

## 1. INTRODUCTION

As the rate-of-living theory suggests, an increase in metabolic activity accelerates the aging process (Pearl, 1928). However, one order of animals defies this general rule – Chiroptera, commonly known as bats. Despite having a high metabolic rate due to energy demanding type of locomotion – active flight – bats exhibit significantly greater longevity than expected for their size and rate of metabolism. Scientists suggest that aging is primarily driven by extrinsic sources of organism’s mortality along with accumulation of toxic byproducts within cells over time (Voigt et al., 2012; Wilkinson and Adams, 2019; Cooper et al., 2024). This raises the question: how do bats overcome these challenges and what mechanisms are responsible for their extended lifespans?

While many studies on bat longevity have predominantly focused on molecular and genetic foundations, relatively few – such as those by Wilkinson and Adams (2019) and Garbino et al. (2021) – have explored the influence of biological traits such as morphology, behavior, and ecology on lifespan (Wilkinson and South, 2002; Healy et al., 2014; Wilkinson and Adams, 2019; Garbino et al., 2021). There remains a notable lack of integrative studies that examine how life-history characteristics contribute to the remarkably low aging rates observed in bats. This gap may, in part, explain the limited availability of high-quality data on bat lifespans.

Body size is most likely to influence bats lifespans, as their type of locomotion determines the survival of individuals and only bats use active flight to move (Voigt et al., 2013; Safi et al., 2013; Sánchez and Carrizo, 2020). We suppose that compact body will lessen use of energy during the flight (Swartz, 1997; Arévalo et al., 2020), at the same time less balanced body size might cause overuse of resources which with time leads to energy deficit and can put individuals in danger of starving or getting ill (extrinsic mortality risks) (Wilkinson and Adams, 2019).

It is commonly assumed that female lifespan may be influenced by maternal investment (i.e., the age at which she reaches maturity, the number of pregnancies throughout her life, the number of offspring per gestation) (Wilkinson and South, 2002). Logically, birth of multiple young increases the energy needed to carry out the pregnancy period and nurture pups until they are old enough to fly and feed themselves (from 6 to 9 weeks) (Garbino et al., 2021). Ultimately, we chose to analyze the reproductive rate in bats, as it most directly reflects the reproductive pressures that may influence female lifespan.

Diet may support the promotion of longevity, especially in bats, which face high mortality risks and must carefully balance energy intake and expenditure (Badal et al., 2020). Bats exploit a wide range of dietary niches, including insects, blood, leaves, nectar, fruit, small mammals (*Megaderma lyra*), or fish (*Myotis vivesi*) (Wilson and Mittermeier, 2019; Cooper, 2024). Corduneanu et al. (2023) have shown the role of diet in shaping the gut microbiome composition, which further influences health and longevity.

Hibernation and torpor have long been argued to enhance survival success in mammals, as both strategies reduce energy expenditure and predation risk through long or short periods of inactivity (Wu and Storey, 2016; Geiser, 2013; Turbill et al., 2016). Early studies in bats hibernation failed to find strong evidence that these forms of torpor significantly prolong the lifespan (Austad and Fischer, 1991). However, more recent findings increasingly support the role of hibernation in promoting longevity in bats (Wilkinson and Adams, 2019; Turbill et al, 2011; Sullivan et al, 2020). Thus, we included both climatic preference (i. e. species prevalent in cold or warm regions) and the presence or absence of hibernation traits possibly influencing bat longevity.

The primary objective of our study was to reconstruct potential patterns in the evolution of longevity in bats and to assess whether the ancestors of modern bats exhibited similarly extended lifespans compared to other mammals. Additionally, we aimed to build upon the work by Healy et al. (2014), Wilkinson and Adams (2019), and Garbino et al. (2021) in identifying the morphometric, behavioral, ecological, and geographical factors that most strongly influence bat lifespan. We suggest that prolonged lifespan is important for bats, as they must mitigate high mortality risks by maintaining a compact body size, compensating for energetic demands of powered flight (Voigt et al, 2012), regulating calorie intake, and surviving the physiologically demanding periods of pregnancy and nursing. Therefore, extended lifespan is essential for animals like bats to reach sexual maturity and reproduce successfully despite previously mentioned extrinsic mortality risks (Wilkinson and Adams, 2019).

To test the hypotheses concerning possible effects on bats lifespan we used correlation and regression analyses. At first stage, we accessed the correlations between the examined traits and evaluated the independent effect of each variable on lifespan. Then the multiple regression model was constructed to reveal the direct contributions of studied variables. Additionally, we identified possible evolutionary patterns for lifespan and morphometric traits.

## METHODS AND MATERIALS

### 1.1. Data collection

We gathered data on maximum lifespan and natural history traits for 108 bat species from open data sources and peer-reviewed studies. All the analyzed bat species come from 11 extant families. The largest part of those species belongs to family Vespertilionidae (55 species), the second half of data consists of families Pteropodidae and Phyllostomidae with 21 and 15 species, respectively. Families Rhinolophidae (4 species), Molossidae (3 species), Emballonuridae (3 species), Megadermatidae (2 species), Miniopteridae (2 species), Mystacinidae (1 species), Hipposideridae (1 species) and Noctilionidae (1 species) together composed the one-fourth of the data (Fig. 1).

**Figure 1.**
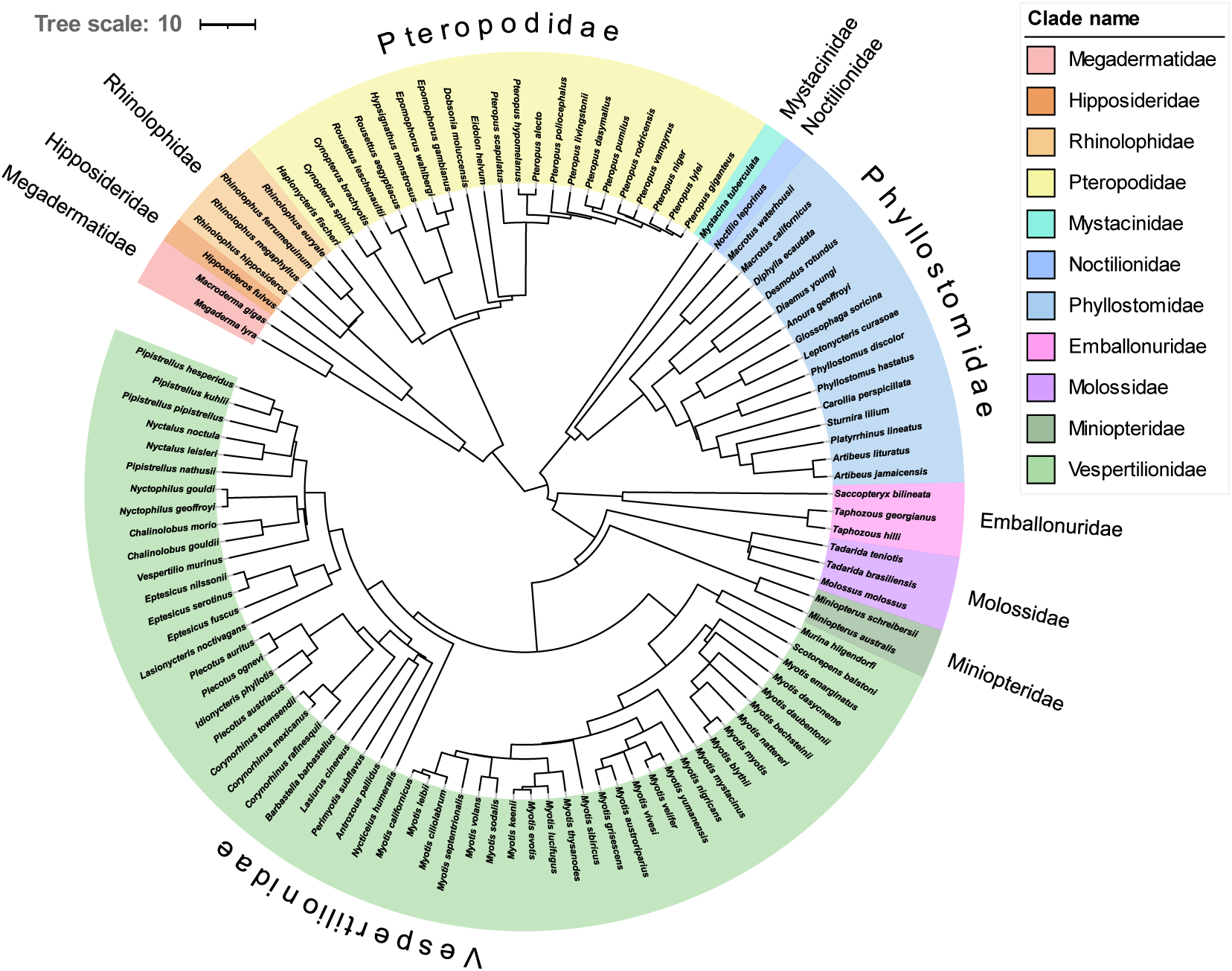
Phylogenetic tree of 108 bat species used in the statistical analysis. Each family is represented by a distinct color to illustrate phylogenetic relatedness (Upham’s et al., 2019; Rothier et al., 2023; Letunic and Bork, 2024).

Life-history traits selected for analysis included morphometrical data (average body length and average forearm length (in mm), average body mass (in g)), longevity (maximum lifespan (in years)), diet (carnivorous, folivores, frugivores, hematophagous, insectivores, nectarivores, omnivorous, palynivores), reproductive pattern (monotocy or polytocy), and climatic preference (prevalence in temperate or tropical regions). Seasonal insectivores from the Phyllostomidae family were marked as omnivores.

In our research, we used taxonomy presented in The Handbook of the Mammals of the World Vol.9 (Wilson and Mittermeier, 2019) and phylogeny from Upham et al. (20199 tree. Information about lifespan was obtained from AnAge database (2023), Wilkinson and Adams (2019) and Garbino’s et al. (2021) original datasets. It is worth mentioning that the main trait in our dataset was longevity. In analysis, we used only the maximum lifespan values because there is not enough information about longevity for most bat species available, or there is only one lifespan estimate available and therefore it was considered as maximal.

The aim of this study was to make the research as comprehensive as possible; therefore, we strived to include chiropterans from all biogeographical realms in our dataset. Climatic preference data were obtained from The Handbook of the Mammals of the World Vol. 9 (Wilson and Mittermeier, 2019) as well as from original articles published in Mammalian Species, particularly those covering North and East American, Western European, and Australo-Asian species. Data on home ranges for Eastern European and Central and East Asian species were gathered from Mezhzherin and Lashkova (2013) (Supplementary Table 1).

To investigate the allometric relationships in bats, we used data for body length, body mass, and forearm length. For statistical analysis, the morphometric data was collected from one single source (Wilson and Mittermeier, 2019) to avoid potential inconsistencies that might arise from using multiple datasets. Additionally, we identified whether there are any associations between longevity and species reproductive pattern, diet, and climatic preference.

For the phylogenetic regression, evolutionary model evaluation, and identification of phylogenetic signal, we used a mammalian phylogenetic tree originally published by Upham et al. (2019) and later adapted by Rothier et al. (2023), which includes a total of 5911 species. The obtained phylogenetic tree occurred to be non-ultrametric for reasons of numerical precision; therefore, we converted it to an ultrametric by force.ultrametric() function from the phytools package (version 2.3-0; Revell, 2024). The final phylogenetic tree used in the analysis, shown on Figure 1, was visualized using the Interactive Tree of Life platform (iTOL v7; Letunic and Bork, 2024).

### 1.2. Statistical analyses

Statistical data analysis was conducted using the R software system (version 4.4.3; R core Team, 2024). Prior to analysis the morphometric data were log-transformed.

To visualize the effects of life-history on maximum lifespan, we first constructed a correlation chart for four variables using the PerformanceAnalytics package (version 2.0.8; Peterson and Carl, 2024) to examine relationships and potential influences of morphometric traits on bat longevity. The chart employs Kendall’s correlation coefficient to assess the pairwise relationships among the variables (Sen, 1968). We then plotted reproductive patterns, climatic preference, hibernation, and dietary preferences using a correlation plot generated with ggplot2() from the tidyverse package (Wickham et al, 2019). This correlation analysis aimed to identify potential relationships between life-history traits, body size, and longevity. The p-values obtained for Kendall’s correlation and phylogenetic signal were adjusted for multiple comparison by p.adjust() function using Benjamini and Hochberg (1995) method for phylogenetic signals and Benjamini and Yekutieli (2001) method for correlation coefficients.

We calculated the phylogenetic signal for maximum lifespan, average body length, body mass and forearm length using Blomberg’s K test (Blomberg et al., 2003) and incorporated sampling error (Ives et al., 2007) with the phylosig() function from the phytools package (version 2.0; Revell, 2024).

In this research, we used the Brownian motion (BM), Ornstein-Uhlenbeck (OU), white noise (WN), and early burst (EB) models as possible explanations for patterns and mechanisms of evolution of bat lifespan, body mass, and forearm length. In short, Brownian motion model assumes random walk of a species mean trait value affected solely by the genetic drift with the correlation structure among trait values proportional to the extent of shared ancestry for pairs of species (Felsenstein, 1973; Lande, 1976). At the same time, the Ornstein–Uhlenbeck model extends the previous model by incorporating the effects of natural stabilizing selection that drive a trait to evolve toward the certain adaptive peak or optimal value (Lande, 1976; Hansen, 1997; Butler and King, 2004). The white noise model, on the other hand, assumes that trait values fluctuate around the optimum with no correlation between pairs of species (Hansen et al., 2008). The last is the early burst model which assumes that the trait accelerates or decelerates exponentially over time (Blomberg et al., 2003; Harmon et al., 2010). To select the models that best fit evolution of longevity and morphometrical data better, we estimated the Akaike Information Criterion (AICc) values for small datasets (Akaike, 1974; Suguira, 1978; Burnham et al., 2011). Subsequently, we applied the fitContinuous() function from geiger package (version 2.0.11; Pennell et al., 2014) to fit the selected models to our comparative data.

Pairwise correlations (see above) provided preliminary information about the possible influences of the studied traits on longevity. But this information can be misleading, since these traits themselves are closely correlated. Therefore, at the next stage of the research we used multiple regression of longevity on the combination of the traits used as predictors. In such an analysis, each regression coefficient best describes the dependence of longevity on the corresponding predictor after removing the effects of other predictors; thus the regression coefficients distinguish the direct effects of the studied traits on longevity from the set of various direct and indirect effects (Lande and Arnold, 1983).

Since phylogenetic signal was confirmed for maximum lifespan in bats (see below the Results section), we applied a phylogenetically corrected regression model using pgls() function from the caper package (version 1.0.3; Orme et al., 2023; Freckleton et al., 2002). To avoid multicollinearity, only one morphometric trait – average forearm length – was used as a predictor. Maximum lifespan (LS) was treated as response variable, while the predictors were: average forearm length (FL, litter size (monotocy vs polytocy, M_vs_P), preferred diet (Diet), climatic preference (species prevalent in cold vs warm regions, FT_vs_SR), hibernation (presence or absence of hibernation, Hibernation). Dietary data were dummy-coded, with insectivory – the dominant dietary type in Chiroptera – set as a reference level (Draper and Smith, 1998). Branch length transformations (lambda, delta and kappa) were optimized to determine the best-fitting model via maximum likelihood transformation.

## 2. RESULTS

### 2.1. Basic pattern

According to the correlation chart (Fig. 2), no significant correlation was found between lifespan and two morphometric traits – body mass (Kendall’s r = 0,114; adjusted p = 0,248) and body length (Kendall’s r = 0,087; adjusted p = 0,460). A very weak correlation was eventually observed between lifespan and forearm length (Kendall’s r = 0,121; adjusted p = 0,246) (Fig. 3a). In contrast, correlations among morphometric traits are very strong: correlations between body mass and forearm length (Kendall’s r = 0,811; adjusted p < 0.001) (Fig. 3b), and between body mass and body length (Kendall’s r = 0,804; adjusted p < 0.001), and between body length and forearm length (Kendall’s r = 0,754; adjusted p < 0.001). Forearm length was selected to represent body size in the multivariate part of our research, while estimating its influence on lifespan, as forearm length best reflects the variation in the other two morphometric traits (Kamran et al, 2013).

**Figure 2.**
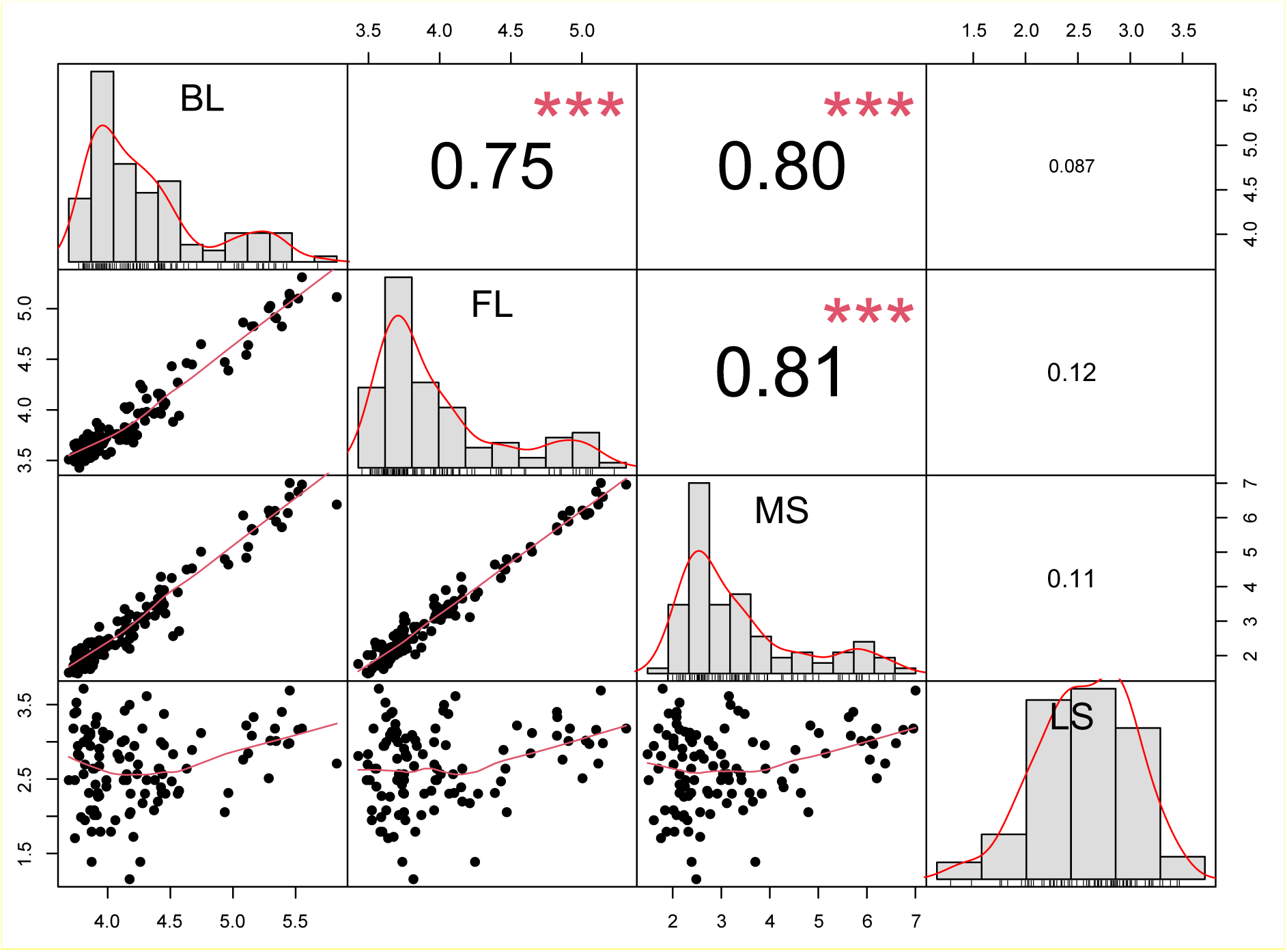
The correlation chart for body length (BL), forearm length (IF), body mass (MS) and lifespan (LS) traits with Kendall’s correlation coefficient.

**Figure 3.**
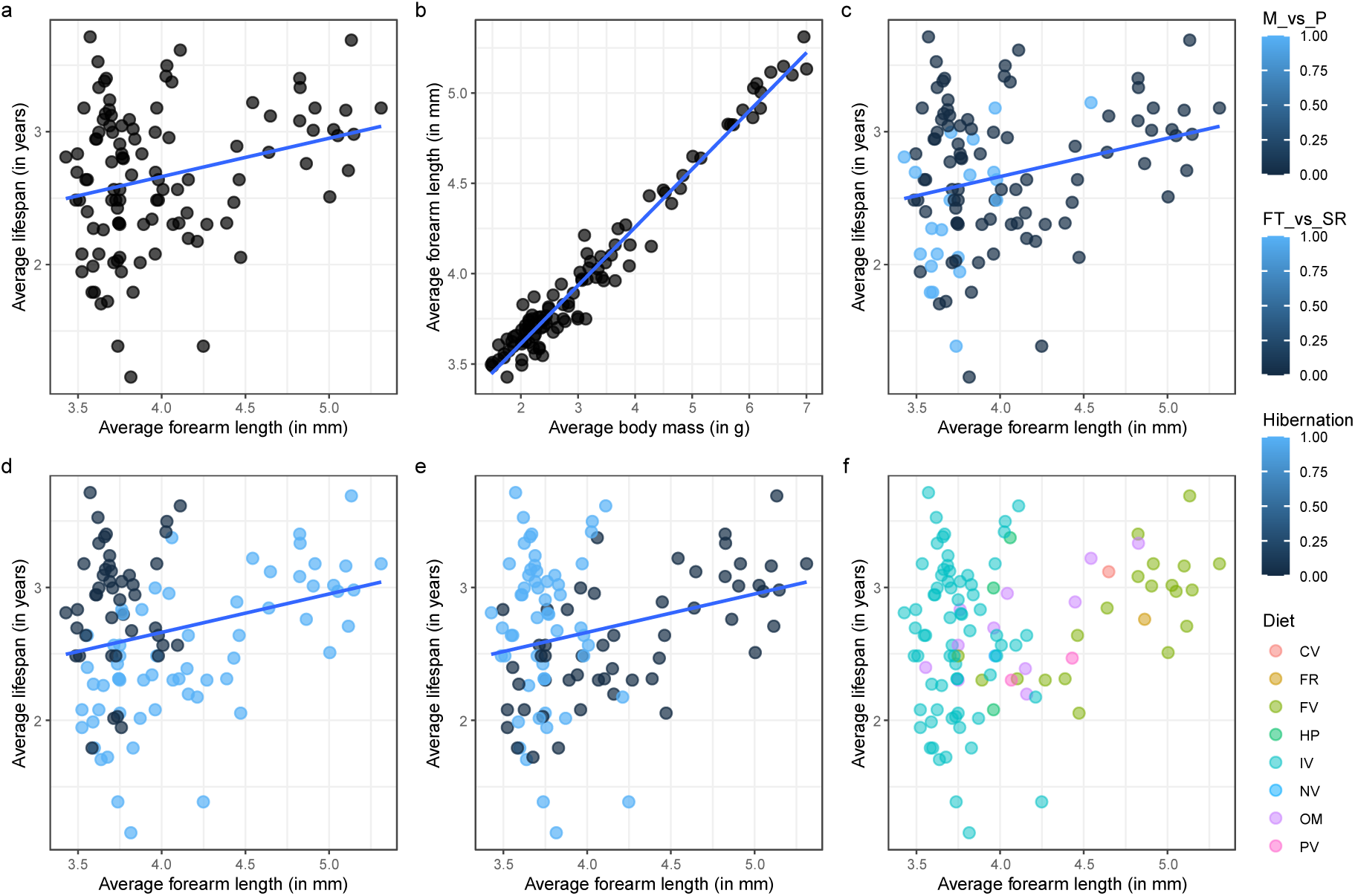
(a) Strong correlation between average body mass and average forearm length. (b) Maximum longevity plotted against average forearm length with overlays of data on (c) reproductive pattern, (d) climatic preference, (e) hibernation and (f) dietary preferences.

Looking at the correlation plot (Fig. 3c), most bat species typically give birth to a single pup per gestation, however, polytocous bats, which can have multiple pups, tend to be smaller in size and have shorter lifespans. We also observed that the bigger part of analyzed bat species that thrive in temperate and cold regions are smaller and likely hibernate or migrate. Tropical bats, on the other hand, display a wider range of body sizes, and probably have diverse adaptations and ecological niches within warmer environments (Fig. 3d). On average, hibernating bats tend to have shorter forearms and generally live longer (Fig. 3e).

In contrast, non-hibernating bats show greater variability in size, with some species possessing significantly longer forearms, including the largest bat species. There is a potential trade-off between forearm length and lifespan that may be associated with hibernation behaviors. Furthermore, correlation plot on Fig. 3f shows that bats with shorter forearms typically have an insect-based diet, whereas those that primarily consume fruits are larger in size. We also observed a very weak negative correlation between Reproductive pattern (monotocy vs polytocy) and Climatic preference (whether species prevalent in cold or warm regions), with Kendall’s r = -0.154. In contrast, the correlation of Climatic preference with Hibernation is very strong (tetrachoric correlation coefficient = -0.86), suggesting an inverse relationship, where species from colder climates are much more likely to hibernate compared to those from warmer regions. Overall, our findings underscore the complex interplay between bat morphology, diet, reproductive strategies, and environmental conditions in shaping a pattern of bat longevity.

Further comparative analysis showed that insectivorous bats are predominant in temperate and cold regions (44 among the studied species), whereas dietary preferences in warm regions are more diverse, including insectivory (25 species), frugivory (20 species), and seasonal omnivory (10). The possible reasons behind the conservative diet in colder regions and greater dietary diversity in warmer regions will be discussed in the Discussion section.

### 2.2. Phylogenetic signal and evolutionary models

A statistically significant phylogenetic signal was detected for both lifespan and morphometric traits (Table 1). Blomberg’s K for maximum lifespan is less than one (K < 1; p < 0.05) indicating that lifespan is relatively conserved across the order Chiroptera, with closely related species exhibiting similar longevity. In contrast, average forearm length and body mass showed high phylogenetic signal values (*K* > 1; *p* < 0.05), suggesting strong phylogenetic clustering and greater trait variability among lineages. Reconstructions of the evolutionary trajectories of these traits are presented in the Supplementary Materials (Figures 4–7).

**Table 1.** Phylogenetic signal estimates and Akaike weights for evolutionary models.

| Traits | Phylogenetic signal |  | Akaike weights for Evolutionary Models |  |  |  |
| --- | --- | --- | --- | --- | --- | --- |
|  | K | Adjusted p-value | BM | OU | EB | WN |
| Maximum lifespan | 0.196 | < 0.001 | < 0.001 | 0.998 | < 0.001 | 0.002 |
| Average forearm length | 1.046 | < 0.001 | 0.057 | 0.924 | 0.020 | < 0.001 |
| Average body mass | 1.006 | < 0.001 | 0.024 | 0.968 | 0.008 | < 0.001 |
| Average body length | 1.032 | < 0.001 | 0.029 | 0.961 | 0.010 | < 0.001 |

\* Abbreviations of statistics and evolutionary models are in the “Materials and Methods” section.

The Akaike weights used to compare the data fit across evolutionary models are also presented in Table 1. In all cases, the Ornstein–Uhlenbeck (OU) model emerged as the best supported model (AICc_LS_ = 0.998, AICc_FL_ = 0.924, AICc_MS_ = 0.968, AICc_BL_ = 0.961). The OU model had a much higher probability than the Brownian motion (BM) and early burst (EB) models, however a strong phylogenetic signal was observed for all morphometric traits (Blomberg’s K > 1), preventing us from rejecting both the BM and EB models in these cases. The results suggest that the dynamics of studied morphometric traits could probably be shaped by a combination of evolution towards a single optimum value – 55.9 mm for forearm length, 25.4 g for body mass, and 73.1 mm for body length – along with random drift. The evolutionary pattern of lifespan, while also best explained by the OU model, differed in important ways. Although a statistically significant but weak phylogenetic signal was detected (K < 1), the OU model had overwhelmingly higher support than competing models — the second-best model (white noise, WN) was 188.2 times less probable. This suggests that lifespan exists under stronger stabilizing selection, evolving toward an adaptive peak estimated at approximately 13.5 years. In contrast to the morphometric traits, lifespan appears less phylogenetically structured and more tightly constrained, likely reflecting selective pressures associated with survival and reproduction strategies in bats.

These findings suggest that while morphometric traits in bats evolve under selection pressure or a combination of drift and selection, lifespan is subject to stronger stabilizing selection, evolving around a distinct adaptive peak with limited influence from phylogenetic relatedness.

### 2.3. Analyzing factors affecting longevity

The phylogenetically informed regression model was used to test the effects of body size, reproductive strategies, environmental factors, and behavioral traits on bat longevity, all of which were hypothesized to have an influence. While the overall model was proved to be statistically significant (F = 7.445, df = 11 and 96, p < 0.0001), the explanatory power was moderate (R² = 0.399): much of the disparity in bat lifespan is caused by unknown factors. Regression coefficients (Table 2) show that maximum lifespan is significantly influenced by average forearm length (as a proxy for body size), litter size, specific diets (hematophagy and omnivory), and climatic preference. Other dietary types and the presence or absence of hibernation had no significant impact. A small yet statistically relevant phylogenetic effect was also detected (Table 1).

**Table 2.** Estimates of the regression coefficients and the intercept in a model of phylogenetically informed regression of average lifespan on a set of morphological, ecological, and life-history predictors.

|  | Estimate | Std. error | t-value | p-value |
| --- | --- | --- | --- | --- |
| Intercept | 0.910 | 0.565 | 1.610 | 0.111 |
| **Forearm length** | **0.518** | **0.148** | **3.503** | **0.001** |
| **Reproduction pattern** | **-0.269** | **0.105** | **-2.566** | **0.012** |
| Frugivorous diet | 0.171 | 0.201 | 0.853 | 0.396 |
| Folivorous diet | 0.048 | 0.454 | 0.106 | 0.916 |
| **Hematophagous diet** | **0.555** | **0.263** | **2.112** | **0.037** |
| Nectarivorous diet | 0.236 | 0.425 | 0.555 | 0.580 |
| **Omnivorous diet** | **0.371** | **0.165** | **2.253** | **0.027** |
| Palynivorous diet | -0.008 | 0.318 | -0.025 | 0.980 |
| Carnivorous diet | 0.517 | 0.444 | 1.166 | 0.247 |
| **Climatic preference** | **-0.719** | **0.108** | **-6.660** | **< 0.001** |
| Hibernation | 0.117 | 0.125 | 0.941 | 0.349 |

\*Significant effects are shown in bold.

## 3. DISCUSSION

Bats are exceptionally long-lived compared to other mammals of similar body size. This longevity is thought to result from a combination of unique life-history traits and molecular mechanisms that mitigate common mortality risks (Healy et al., 2024; Gorbunova et al., 2020; Wilkinson and Adams, 2019; Garbino et al., 2021; Cooper et al., 2024). Recent studies suggest complex interrelations among morphometric variables, life-history traits, and longevity in bats. Lifespan extremes are influenced not only by body size but also by factors such as forearm length (a proxy for body size), reproductive strategies, environmental conditions, and dietary preferences. Our findings generally align with these conclusions, supported by our regression results (Wilkinson and Adams, 2019; Garbino et al., 2019).

Bats exhibit high average and maximum lifespans. For example, the generation length for *Myotis lucifugus* (Vespertilionidae) was estimated to be 4.76 years, at the same time life expectancy at birth – 2.85 years, and life expectancy at maturity – 5.90 years (Humphrey and Cope, 1976; Heppell et al., 2000). Similarly, based on the data collected by Fleming (1988, quoted in Heppell et al., 2000), Heppell et al. (2000) estimated generation length for *Carollia perspicillata* (Phyllostomidae) is 3.14 years with life expectancy at birth and maturity being 2.56 years and 2.90 years, respectively. From the demographic data collected by Horáček et al. (1984), the generation length for *Myotis myotis* (Vespertilionidae) was estimated to be 4.4 years (Dzeverin, 2008). For *Myotis bechsteinii*, generation time was estimated to be about two to three years (Kerth et al., 2003): the considerable difference with the estimates obtained for two other *Myotis* species can probably be due to the different estimation techniques.

Body size is a fundamental factor influencing nearly every aspect of an animal’s biology, from development and metabolism to ecological interactions and behavior. In comparative studies, body mass is commonly used to assess correlations between morphology and function (Hubel et al., 2010). In bats, however, the forelimbs play a particularly critical role. It is also worth mentioning that in numerous studies, forearm length has been recognized as a more reliably measured proxy for body size than body mass, as the mass tends to fluctuate throughout the day or across seasons (Thiagavel et al. 2017). These complex and highly specialized structures — composed of membranes, muscles, bones, and joints — are essential for powered flight, the defining characteristic of the order Chiroptera, whose name literally means “hand-wing”. Among the forelimb components, the forearm is especially important for supporting wing membranes, contributing to wing mass, and maintaining structural integrity during flight (Sánchez and Carrizo, 2020). Due to its structural and functional relevance, we selected average forearm length as the primary morphometric trait in our regression analysis, rather than body mass or body length. Although the Ornstein–Uhlenbeck (OU) model identified a single optimum forearm length of 55.9 mm for the order, this should not be interpreted to mean that only small-bodied bats can achieve exceptional longevity. Meanwhile, our evolutionary model analyses suggest the existence of multiple (at least, two) optima for forearm length, body length, and body mass.

Testing for phylogenetic signals revealed that lifespan is consistently high across Chiroptera, suggesting that extended longevity has been an evolutionary characteristic of bats for a long time — likely exceeding that of similarly sized mammals. Our findings for body mass phylogenetic signal align with previous research that reported a strong phylogenetic signal for this trait in bats (Safi et al., 2013). Similarly, Wilkinson and Adams (2019) found a high Pagel’s λ value for body mass (λ = 0.88), indicating trait similarity consistent with a Brownian motion (BM) model of evolution. In our study, we also detected a strong phylogenetic signal for body length and forearm length. These morphometric traits were expected to vary among clades in a manner expected under Brownian motion, the similar evolutionary patterns accord with the strong correlations observed among body mass, body length, and forearm length. However, model fitting results suggest that the evolution of these traits is better described by the Ornstein–Uhlenbeck (OU) model, which assumes evolution toward a stabilizing optimum rather than random drift. According to the Akaike Information Criterion corrected for small sample size (AICc), the evolution of lifespan in Chiroptera is best explained by the OU model (with the WN model in second place with the large gap from the former model). This indicates that bat lifespan is evolutionarily regulated by selection, as opposed to evolving randomly under a Brownian motion model. In contrast, the evolutionary patterns of forearm length and body mass show high support for the OU, whereas Brownian motion (BM) and early burst (EB) models were less supported by our data, but not so much that they can be neglected. Such patterns suggest more complex evolutionary dynamics that shaped these traits. Furthermore, while strong correlations exist among morphological traits, the correlation between these traits and longevity is weak or negligible in our data. This contrasts with previous studies that identified a strong correlation between lifespan and body mass (Cooper, 2024). This discrepancy may result from differences in raw data and research objectives; our focus was to detect general correlations suitable for regression analysis. The observed phylogenetic signal for lifespan also supports the idea that longevity is a conservative trait, implying that ancestral bats likely had longer lifespans relative to other mammals. Conversely, traits such as forearm length and body mass exhibit greater variability, reflecting adaptations to diverse ecological niches and life-history strategies among bat species.

Our study revealed that bat species which hibernate are predominantly insectivorous, tend to have smaller bodies, and are mostly found in colder regions with seasonal temperature variation. Firstly, for many bat groups, insects provide a critical source of energy necessary for powered flight and serve as a valuable protein supplement during the breeding season (Vafidis, 2019). For example, *Myotis lucifugus* can consume up to 1,200 mosquitoes in just one hour (Duccummon, 2000). Similarly, a colony of 150 big brown bats (*Eptesicus fuscus*) has been recorded consuming approximately 123,000 insects such as Cicadellidae, *Diabrotica*, *Acalymma*, *Phyllophaga*, and Pentatomidae during a single breeding season (Whitaker, 1995). The caloric content of the diet, the protein-to-carbohydrates ratio, and the energy costs associated with foraging or hunting directly impact bats’ exposure to predators (Couteur et al., 2015; Wilkinson and South, 2002). Secondly, bats in temperate regions must expend significant energy to maintain body temperature, as ambient temperatures are much lower than in tropical regions. Thus, an insectivorous diet is particularly advantageous in colder climates due to the high energy yield it provides (Vafidis, 2019). When unfavorable environmental conditions arise—such as subzero temperatures or prolonged wet and stormy weather — bats enter brief periods of torpor or extended hibernation to reduce energy expenditure (Geiser and Turbill, 2009). Hibernation is more common among bat species inhabiting temperate zones and higher latitudes with seasonal temperature changes (Wilkinson and South, 2019; Geiser, 2013). Howevert – there are exceptions – such as the species *Cynopterus brachyotis*, which exhibits heterothermy despite living in warm tropical climate (Welman, 2017).

All bats from temperate regions are heterothermic, and their body size appears to be evolutionarily shaped by environmental conditions, especially ambient temperature (Geiser, 2013). In mammals, the surface-area-to-volume ratio is important in determining heat production and dissipation rates (Bergmann, 1998). Bergmann’s rule predicts that endotherms tend to evolve larger body sizes in colder environments to conserve heat. However, bats appear to deviate from this pattern; our findings show that bat species prevalent in temperate regions are generally smaller, while the largest bat species are found in tropical and subtropical regions. This pattern is supported by other studies, which note that insectivorous bats often deviate from Bergmann’s rule (Meiri, 2003). A study on Neotropical frugivorous bats examined Bergmann’s rule through the lens of heat transfer. It found that smaller species compensate for low temperatures by increasing their body size, whereas larger species increase fur density to enhance thermal insulation in cold environments (Castillo-Figueroa, 2022). In contrast, Freckleton (2003) argued that larger species are more likely to follow Bergmann’s rule than smaller ones. Hibernation and torpor serve as mechanisms for bats to avoid starvation during periods of scarce food and water supplies. The accompanying metabolic slowdown causes body temperature to drop, making hibernating bats less detectable to predators and increasing survival chances (Wu, 2016). While hibernation is generally believed to positively influence bat longevity by conserving energy, this relationship remains debated. For instance, Brunet-Rossini (2004) found no significant difference in lifespan between hibernating and non-hibernating bat species. Conversely, a more recent study by Wilkinson and Adams (2019) showed a correlation between hibernation and increased lifespan, supporting the idea that bats in temperate zones may live longer due to extended periods of metabolic stasis. Additional recent studies further reinforce the role of hibernation in promoting longevity (Wilkinson and South, 2002; Turbill et al., 2011; Sullivan et al., 2020). However, our own data demonstrate no significant correlation between hibernation and lifespan in bats, further reinforcing the uncertainty surrounding the role of this trait in influencing longevity.

As previously mentioned, most bat species give birth to a single offspring per gestation (monotocous), while only a few are polytocous, producing two or more pups (Garbino et al., 2021). Analysing reproductive strategies, we observed that monotocous bats span a wide range of dietary preferences — from carnivory to herbivory — whereas polytocous species are predominantly insectivorous. This pattern may reflect the higher energetic demands of multiple offspring, making an insect-based diet more favorable. Additionally, monotocous bats are distributed across both warm and cold climates, while polytocous species are largely confined to colder regions. We hypothesized that reproductive strategies in bats might be shaped by dietary and environmental factors — consistent with previous research by Wilkinson and Adams (2019), which associated polytocy with hibernation and colder climates. However, our analysis did not reveal significant support for the hypothesis that hibernation affects bat longevity.

In this research, we did not analyze bat roost types nor group size as potential traits influencing bat lifespan. These two factors have been examined and described in detail in the work by Garbino et al. (2021). Although these variables were not included in our analysis, a review of the existing literature, including the findings from Garbino et al. (2021), show that group size does not have a substantial effect on bat longevity.

In most bat species, females aggregate at communal roosts during breeding and nursing seasons This social structure generally reduces competition for resources, and while occasional aggression may occur, it is considered rare (Lukas and Clutton-Brock, 2020). Reproduction is itself quite demanding process, especially in the order Chiroptera, as the size of baby is quite high compared to other mammals, for example, average body mass for *Pipistrellus subflavus* (or *Perimyotis subflavus* according to modern classification) females is 5.8 g and body mass of a pup on average is 1,8 g (Hoying and Kunz, 1998). Additionally, the high-maintenance lifestyle of bats makes many life aspects more complicated and energy-demanding. Such drastic increase in body size in pregnant females increases extrinsic mortality rates and pregnancy itself stresses out the cells. Therefore, reproductive factors such as the number of pups per gestation, are important in determining bat lifespan. Though polyembryony sometimes occurs as an individual abnormality even in obligately monotocous bat species, selection seems to act strongly against such variations (Arlettaz, 1993). High risks associated with pregnancy and birthing in bats increase female mortality, especially in species that reach sexual maturity early and produce multiple offspring per litter. Our findings are consistent with previous research, which has shown that bat lifespan is closely associated with age at sexual maturation and overall reproductive investment. Females that mature earlier and are polytocous (bat species that give one pup per gestation) have higher chance of mortality at both the physiological and molecular levels due to stress and risks associated with frequent reproduction. Thus, females with a faster reproductive rate and higher reproductive output are expected to have shorter lifespans, emphasizing the trade-off between reproductive effort and longevity (Lagunas-Raguel, 2019). Due to the adaptation to powered flight, low fecundity and relatively very large embryos are the fixed constant features of bats. Increasing lifespan seems to be a mechanism evolved to compensate for these constraints.

The bats of most families can only live in regions with tropical and subtropical climates. Vespertilionidae is the only bat family whose clades and lineages have successfully colonized regions with temperate and even relatively cold climates in Palearctics and Nearctics. This was achieved by changes in reproductive strategy, rearrangement of reproductive organs, the emergence of the ability to polyembryony (Borissenko, 1999, 2000), as well as complex ecological, ethological, and physiological adaptations such as migration and hibernation.

Regarding reproductive strategies: monotocous bat species exhibit a wide range of dietary habits, ranging from carnivory to herbivory, whereas polytocous bats are predominantly insectivorous. We suggest that this pattern may result from the higher energetic demands faced by polytocous females during pregnancy, making an insect-based diet (rich in protein and energy) more effective. Additionally, our data suggests that monotocous species are distributed across both warm and cold regions, while polytocous bats are more commonly found in colder environments. We tested whether reproductive strategies in bats may be influenced either by diet or environmental conditions, or these factors both, with polytocous bats likely hibernating and adapting to colder climates – where their higher reproductive output and insectivorous diet may help meet increased energy demands.

Interestingly, one of the most unexpected findings of our study is that only certain dietary types appear to positively influence bat longevity. Specifically, this effect was observed in hematophagous species, seasonal insectivores, and fully omnivorous bats. We presume that these diets may have some survival advantages, potentially through enhanced adaptations to changing environments, thereby reducing physiological stress from periods of starvation, or underlying molecular mechanisms. Nonetheless, we found little direct support for this idea in the existing literature, except for Wilkinson and South (2002), who reported no association between diet and longevity at all.

A major challenge we encountered during data collection was the lack of longevity records for many bat species. While species native to North American and Europe are relatively well-documented, those species inhabiting tropical and subtropical regions are heavily underrepresented in the literature. For instance, despite there being over 1 400 known bat species (Wilson and Mittermeier, 2019), our dataset includes information on only 108. Additionally, instead of using average lifespan values we were compelled to rely on maximum lifespan data, as most species had only a single record of longevity. This finding underscores that more systematic research on bat lifespan is needed to enable a comprehensive understanding of their exceptional longevity.

Our findings provide a new framework for understanding bat longevity and may contribute to future research on the evolution of life-history traits in mammals.

## CONCLUSION

Whereas forearm length, body mass and body length vary more strongly with phylogeny, lifespan in bats shows a moderate phylogenetic signal and is relatively conserved across species. Data analysis revealed that bat lifespan is shaped by forearm length (used as a proxy for general body size), reproduction rate, certain dietary types (hematophagy and omnivory), and climatic preference, while other traits showed no significant effects. Evolutionary model testing for forearm length, body length, body mass, and lifespan indicated that all traits are best described by the Ornstein–Uhlenbeck model, suggesting stabilizing selection toward optimal values. Lifespan emerged as a particularly constrained trait, with an estimated evolutionary optimum of approximately 13.5 years across the order Chiroptera, with some possible variation. These findings underscore the importance morphological and life-history traits in shaping bat longevity.

## ACKNOWLEDGEMENTS

We would like to express our sincere gratitude to Maria Ghazali, Pavel Gol’din, Lena Godlevska, Oleksandr Zinenko, and Oksana Shatkovska for their constructive criticism and invaluable insights. We are also deeply indebted to the staff of the Department of Evolutionary Morphology at the Schmalhausen Institute of Zoology for their continuous support throughout our research. Finally, our special thanks go out to the Armed Forces of Ukraine, whose efforts have ensured our safety and allowed this work to continue.

## Supplementary Materials

**Figure 4.**
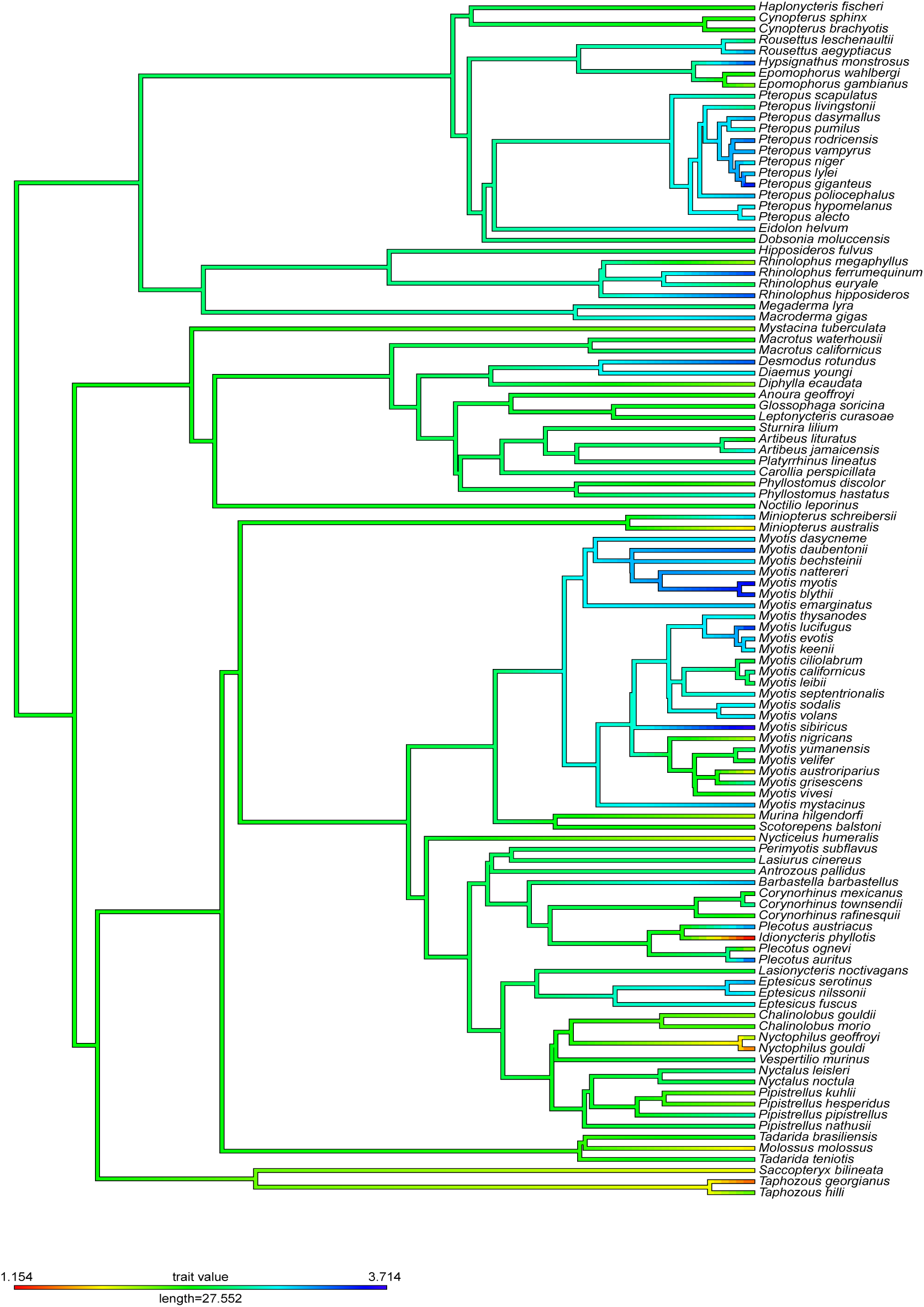
Evolutionary mapping of lifespan on the bat phylogenetic tree (Upham, 2019; Rothier, 2023).

**Figure 5.**
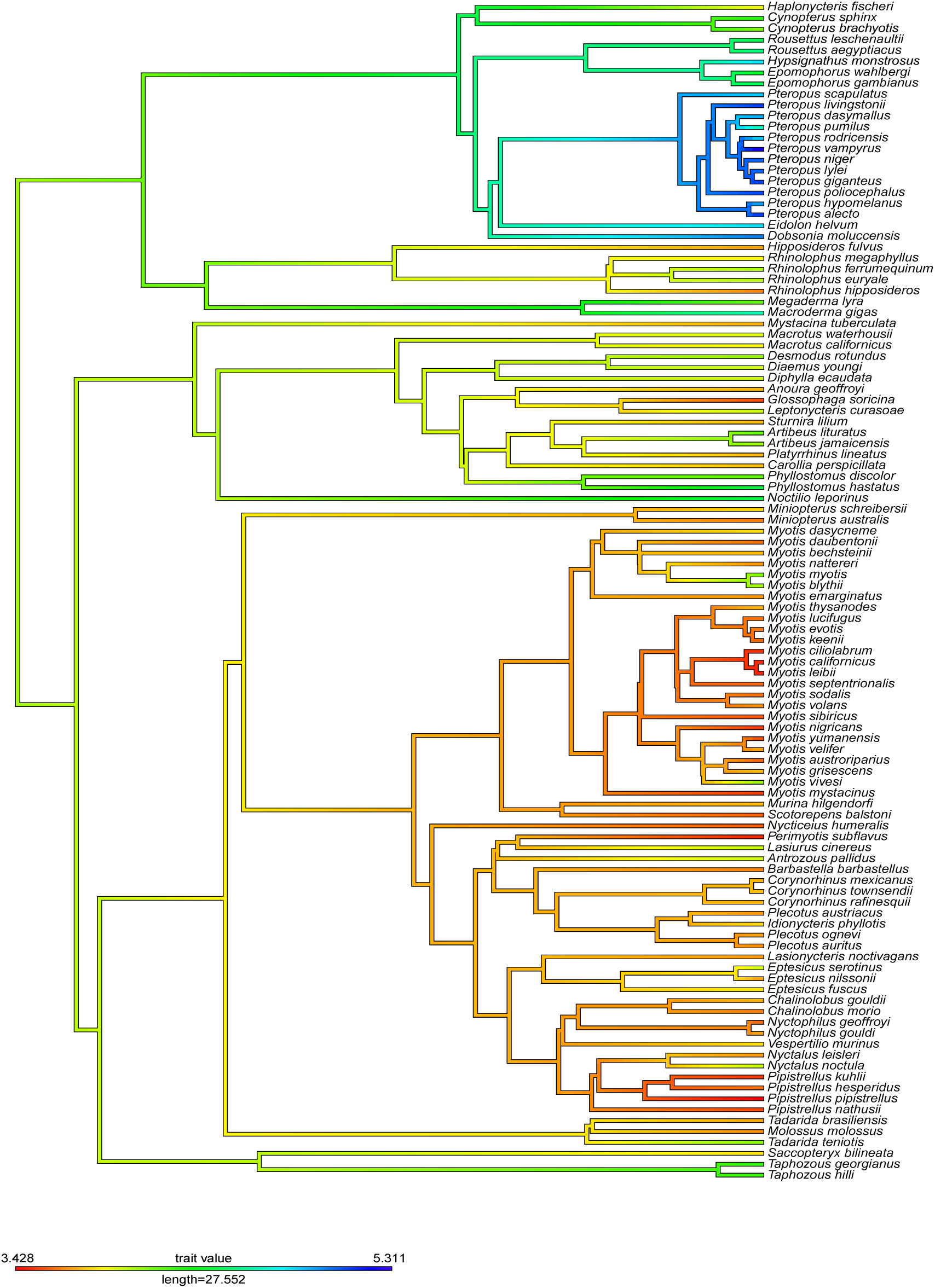
Evolutionary mapping of average forearm length on the bat phylogenetic tree (Upham, 2019; Rothier, 2023).

**Figure 6.**
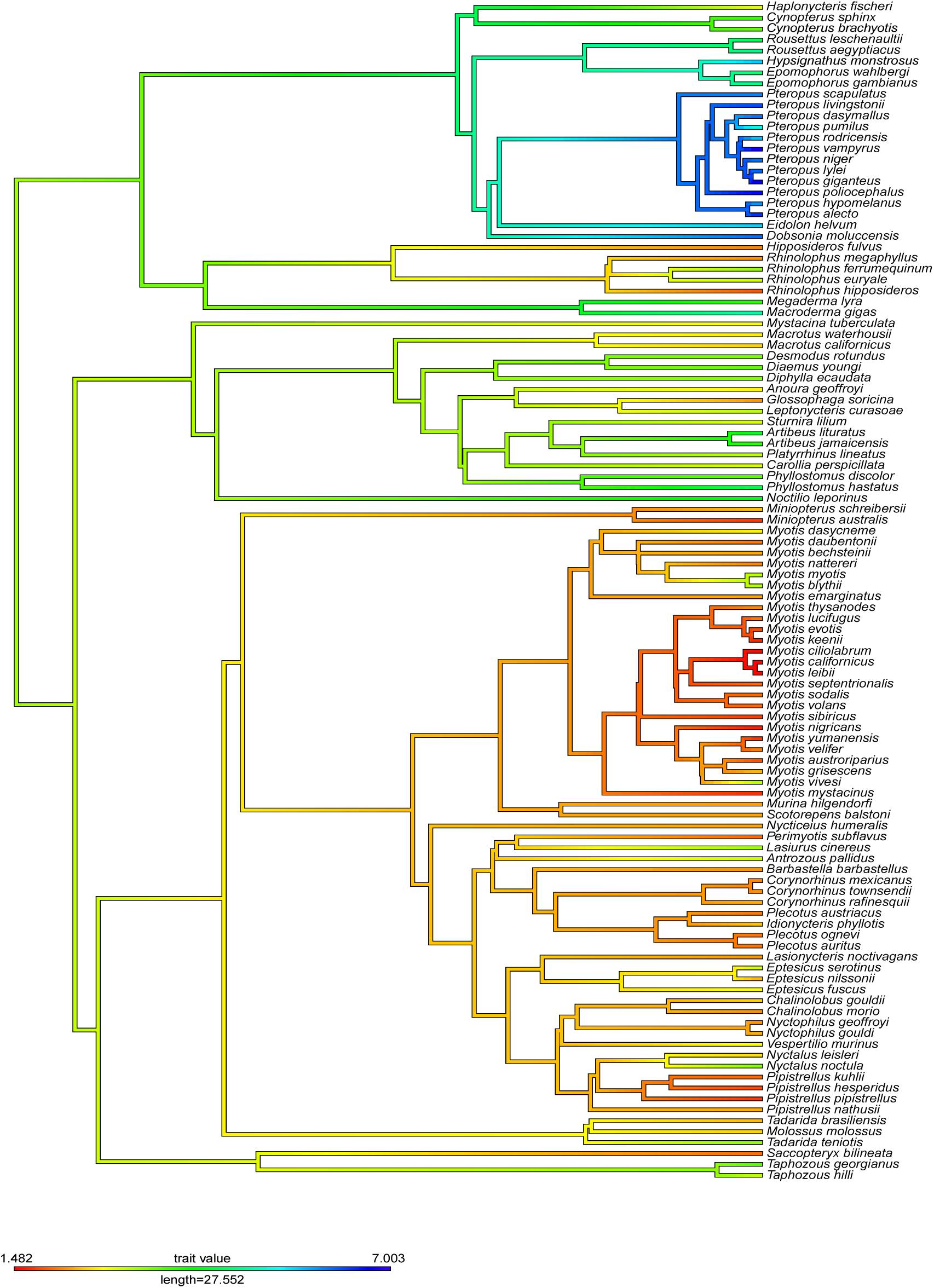
Evolutionary mapping of average body mass on the bat phylogenetic tree (Upham, 2019; Rothier, 2023).

**Figure 7.**
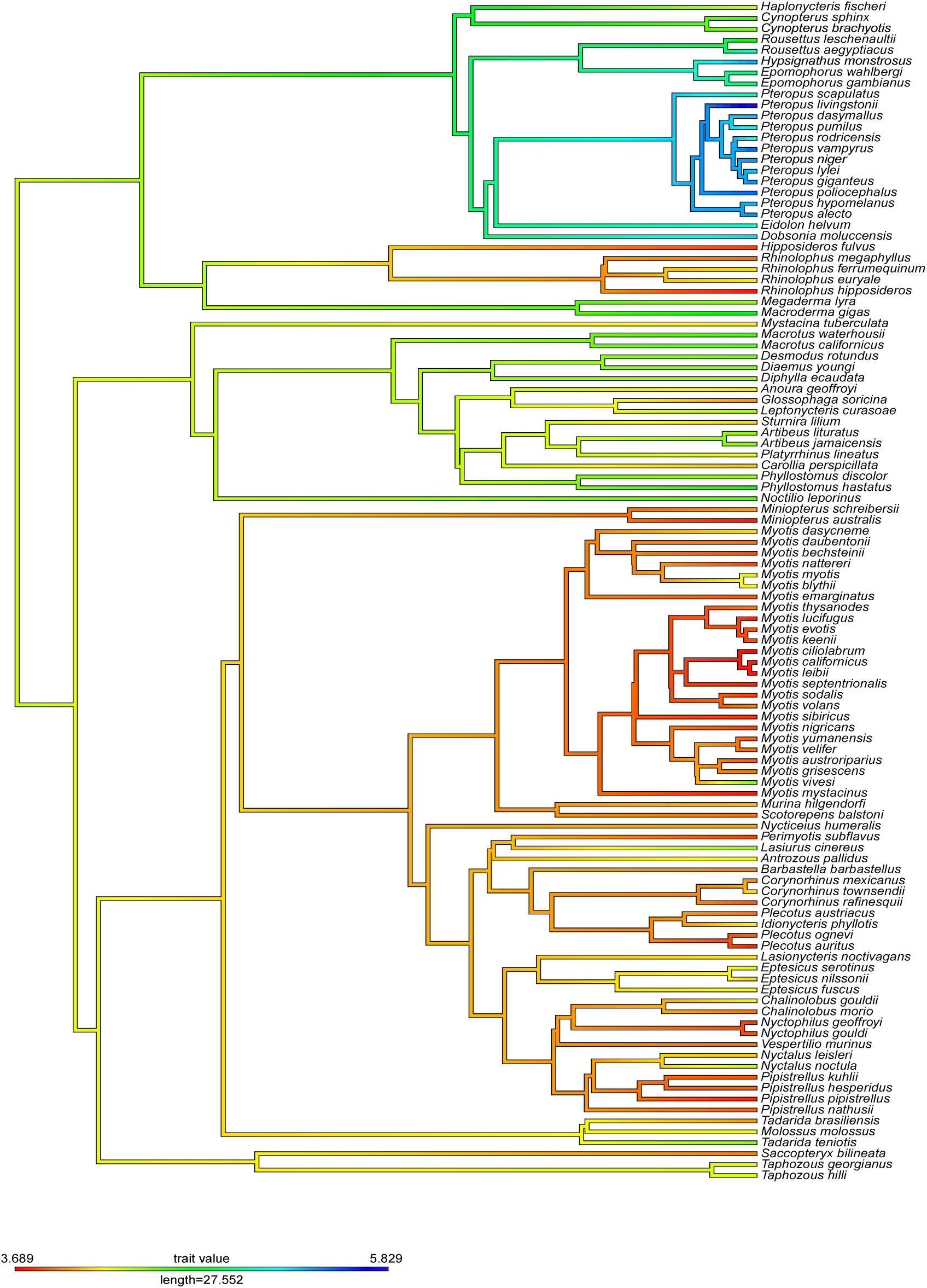
Evolutionary mapping of average body length on the bat phylogenetic tree (Upham, 2019; Rothier, 2023).

| Species name | BL | FL | MS | LS | M vs P | Diet | FT vs SR | Hibernation | Sources | Notes |
| --- | --- | --- | --- | --- | --- | --- | --- | --- | --- | --- |
| *Anoura geoffroyi* | 65,5 | 42,5 | 15,5 | 10 | 0 | OM | 1 | 0 | 5,11,13,21,72 |  |
| *Antrozous pallidus* | 69,5 | 52,6 | 21,5 | 15 | 1 | OM | 0 | 1 | 13,21 |  |
| *Artibeus jamaicensis* | 85,5 | 57 | 49,3 | 19 | 0 | OM | 1 | 0 | 5,11,13,21,72,75 |  |
| *Artibeus lituratus* | 83,5 | 63,5 | 72,5 | 11 | 0 | OM | 1 | 0 | 13,21 |  |
| *Barbastella barbastellus* | 52,5 | 39 | 9,5 | 23 | 0 | IV | 0 | 1 | 5,13,21,23,72,76 |  |
| *Carollia perspicillata* | 59 | 43 | 20 | 17 | 0 | OM | 1 | 0 | 5,11,13,21,72,77 |  |
| *Chalinolobus gouldii* | 65,5 | 42,5 | 13 | 7,8 | 1 | IV | 1 | 1 | 5,11,13,21,78 |  |
| *Chalinolobus morio* | 50,5 | 38,5 | 9,3 | 9,6 | 1 | IV | 1 | 1 | 5,11,13,21 |  |
| *Corynorhinus mexicanus* | 50,5 | 42,3 | 8,5 | 10 | 0 | IV | 1 | 1 | 13,21 |  |
| *Corynorhinus rafinesquii* | 47 | 42,2 | 10,8 | 10 | 0 | IV | 1 | 1 | 13,21 |  |
| *Corynorhinus townsendii* | 65 | 43,4 | 9 | 16,4 | 0 | IV | 0 | 1 | 13,21,72 |  |
| *Cynopterus brachyotis* | 84 | 60,5 | 36 | 10 | 0 | FR | 1 | 0 | 13,21,72 |  |
| *Cynopterus sphinx* | 95,5 | 71,5 | 46,3 | 10 | 0 | FV | 1 | 0 | 11,13,21,72,79 |  |
| *Desmodus rotundus* | 85,5 | 58 | 32,5 | 29 | 0 | HP | 1 | 0 | 5,11,13,21,72,80 |  |
| *Diaemus youngi* | 83,5 | 52,5 | 38,5 | 20 | 0 | HP | 1 | 0 | 5,11,13,21,23,81 |  |
| *Diphylla ecaudata* | 79 | 52,5 | 31,5 | 8 | 0 | HP | 1 | 0 | 5,13,21,23,72,82 |  |
| *Dobsonia moluccensis* | 198 | 149 | 495 | 12 | 0 | FR | 1 | 0 | 11,13,21,72 |  |
| *Eidolon helvum* | 173 | 125 | 290 | 22 | 0 | FV | 1 | 0 | 5,11,13,21,72,83 |  |
| *Epomophorus gambianus* | 139 | 87,5 | 121 | 7,8 | 0 | FV | 1 | 0 | 5,11,13,21,72,84 |  |
| *Epomophorus wahlbergi* | 143 | 80,5 | 104 | 10 | 0 | FR | 1 | 0 | 11,13,21,72 |  |
| *Eptesicus fuscus* | 67,5 | 46,5 | 17 | 19 | 1 | IV | 0 | 1 | 5,11,13,21,72,85 |  |
| *Eptesicus nilssonii* | 61 | 40,7 | 11 | 20 | 1 | IV | 0 | 1 | 5,11,13,21,72,86 |  |
| *Eptesicus serotinus* | 72 | 53 | 21,5 | 24 | 1 | IV | 0 | 1 | 5,11,13,21 |  |
| *Glossophaga soricina* | 54 | 35 | 9,5 | 11 | 0 | NV | 1 | 0 | 5,11,13,21,72,87 |  |
| *Haplonycteris fischeri* | 73,5 | 49 | 18,5 | 10 | 0 | FR | 1 | 0 | 11,13,21 |  |
| *Hipposideros fulvus* | 45 | 41 | 8,5 | 13 | 0 | IV | 1 | 0 | 11,13,21,72 |  |
| *Hypsognathus monstrosus* | 219 | 125 | 305 | 30 | 0 | FV | 1 | 0 | 5,11,13,21,88 |  |
| *Idionycteris phyllotis* | 65 | 45,4 | 12 | 3,2 | 0 | IV | 1 | 1 | 11,13,21 |  |
| *Lasionycteris noctivagans* | 63,5 | 40,5 | 9,6 | 12 | 1 | IV | 0 | 1 | 11,13,21,72 |  |
| *Lasiurus cinereus* | 82 | 53,5 | 27,5 | 14 | 1 | IV | 0 | 1 | 11,13,21,72 |  |
| *Leptonycteris curasoae* | 79,5 | 53 | 23,5 | 12 | 0 | NV | 1 | 0 | 13, 21 |  |
| *Macroderma gigas* | 115 | 105 | 150 | 23 | 0 | CV | 1 | 0 | 5,11,13,21,72,89 |  |
| *Macrotus californicus* | 92 | 48,5 | 13 | 17 | 0 | IV | 1 | 0 | 11,13,21 |  |
| *Macrotus waterhousii* | 96,5 | 51,5 | 15 | 10 | 0 | IV | 1 | 0 | 11,13,21 |  |
| *Megaderma lyra* | 82,5 | 64 | 50 | 14 | 0 | IV | 1 | 0 | 11,13,21,72 |  |
| *Miniopterus australis* | 42 | 38 | 5,8 | 5,5 | 0 | IV | 1 | 1 | 11,13,21 |  |
| *Miniopterus schreibersii* | 55 | 45 | 12 | 22 | 0 | IV | 0 | 1 | 5,11,13,21,72 |  |
| *Molossus molossus* | 67 | 39,5 | 13 | 5,6 | 0 | IV | 1 | 0 | 11,13,21,72 |  |
| *Murina hilgendorfi* | 58 | 42,9 | 10 | 7 | 1 | IV | 0 | 1 | 5,11,13,21 |  |
| *Myotis austroriparius* | 48 | 36,5 | 6,5 | 6 | 1 | IV | 1 | 1 | 13, 21, 23 |  |
| *Myotis bechsteinii* | 46,5 | 43 | 10,5 | 21 | 0 | IV | 0 | 1 | 5, 13, 21, 23,72 |  |
| *Myotis blythii* | 65 | 56,3 | 24,3 | 33 | 0 | IV | 0 | 1 | 5, 13, 21, 23,72 |  |
| *Myotis sibiricus* | 45 | 35,6 | 6 | 41 | 0 | IV | 0 | 1 | 5, 13, 21, 23,72 | Known as *Myotis brandtii* in older classification |
| *Myotis californicus* | 43 | 33 | 4,4 | 17 | 0 | IV | 0 | 0 | 13, 21, 23,72 |  |
| *Myotis ciliolabrum* | 40 | 33,4 | 4,5 | 12 | 0 | IV | 0 | 1 | 13, 21, 23 |  |
| *Myotis dasycneme* | 62,5 | 46 | 15,5 | 21 | 0 | IV | 0 | 1 | 5, 13, 21, 23,72,90 |  |
| *Myotis daubentonii* | 50 | 37,5 | 8 | 28 | 0 | IV | 0 | 1 | 5, 13, 21, 23,72 |  |
| *Myotis emarginatus* | 47,5 | 40,4 | 10,5 | 23 | 0 | IV | 0 | 1 | 5, 13, 21, 23,72,91 |  |
| *Myotis evotis* | 47,5 | 38,5 | 6,5 | 22 | 0 | IV | 0 | 1 | 5, 13, 21, 23,72,92 |  |
| *Myotis grisescens* | 51 | 43,2 | 11,5 | 17 | 0 | IV | 1 | 1 | 5, 13, 21, 23,72 |  |
| *Myotis keenii* | 47,5 | 36,8 | 5 | 19 | 0 | IV | 0 | 1 | 13,21,23,93 |  |
| *Myotis leibii* | 42 | 32,7 | 4,5 | 12 | 0 | IV | 0 | 1 | 5, 13, 21, 23,72 |  |
| *Myotis lucifugus* | 42,5 | 37,3 | 8,5 | 34 | 0 | IV | 0 | 1 | 13, 21, 23, 72 |  |
| *Myotis myotis* | 74,5 | 61 | 23,5 | 37 | 0 | IV | 0 | 1 | 5, 13, 21, 23,72 |  |
| *Myotis mystacinus* | 41,5 | 34,3 | 5,5 | 24 | 0 | IV | 0 | 1 | 5, 13, 21, 23 |  |
| *Myotis nattereri* | 45,5 | 39,9 | 8,5 | 24 | 0 | IV | 0 | 1 | 5, 13, 21, 23,72 |  |
| *Myotis nigricans* | 45,5 | 33,9 | 5 | 7 | 0 | IV | 1 | 0 | 13, 21, 23, 72 |  |
| *Myotis septentrionalis* | 43 | 37 | 7,5 | 19 | 0 | IV | 0 | 1 | 13, 21, 23 |  |
| *Myotis sodalis* | 45 | 37,5 | 7,5 | 20 | 0 | IV | 0 | 1 | 13, 21, 23,72 |  |
| *Myotis thysanodes* | 51 | 42,5 | 9 | 18 | 0 | IV | 0 | 1 | 13, 21, 23,72 |  |
| *Myotis velifer* | 49,5 | 41,9 | 8,5 | 11 | 0 | IV | 1 | 1 | 5, 13, 21, 23,72,94 |  |
| *Myotis vivesi* | 86,5 | 58,4 | 25 | 10 | 0 | PV | 1 | 0 | 13, 21, 23,72 |  |
| *Myotis volans* | 53 | 40 | 7,5 | 21 | 0 | IV | 0 | 1 | 13, 21, 23,72 |  |
| *Myotis yumanensis* | 48,5 | 35 | 5,5 | 14 | 0 | IV | 1 | 1 | 5, 13, 21, 23,72,95 |  |
| *Mystacina tuberculata* | 65 | 41,9 | 16 | 7,6 | 0 | IV | 0 | 0 | 5, 13, 21, 23,72 |  |
| *Noctilio leporinus* | 91 | 84 | 70 | 12 | 0 | PV | 1 | 0 | 13, 21, 23,72 |  |
| *Nyctalus leisleri* | 60 | 40,5 | 14 | 16 | 0 | IV | 0 | 1 | 5, 13, 21, 23,72 |  |
| *Nyctalus noctula* | 74,5 | 53,5 | 30,5 | 12 | 1 | IV | 0 | 1 | 5, 13, 21, 23,72 |  |
| *Nycticeius humeralis* | 56 | 36 | 10,2 | 6 | 1 | IV | 0 | 0 | 11,13,21,72 |  |
| *Nyctophilus geoffroyi* | 44,1 | 36,2 | 9,2 | 7,3 | 1 | IV | 1 | 1 | 11,13,21 |  |
| *Nyctophilus gouldi* | 48 | 42 | 10,9 | 4 | 1 | IV | 1 | 1 | 11,13,21 |  |
| *Perimyotis subflavus* | 45,5 | 32,9 | 7,5 | 15 | 1 | IV | 1 | 0 | 5,11,13,21,72 | Known as *Pipistrellus subflavus* in older classification |
| *Phyllostomus discolor* | 81,5 | 64 | 38,5 | 9 | 0 | FV | 1 | 0 | 5,11,13,21,72,96 |  |
| *Phyllostomus hastatus* | 107 | 85,5 | 92,5 | 18 | 0 | FV | 1 | 0 | 13, 21, 23, 72,97 |  |
| *Pipistrellus hesperidus* | 49 | 37,5 | 6,3 | 8 | 1 | IV | 1 | 0 | 13, 21, 23 |  |
| *Pipistrellus kuhlii* | 47,5 | 33,9 | 7,5 | 8 | 1 | IV | 0 | 1 | 5,11,13,21,72 |  |
| *Pipistrellus nathusii* | 49 | 34,7 | 10,8 | 14 | 0 | IV | 0 | 1 | 5,11,13,21 |  |
| *Pipistrellus pipistrellus* | 43,5 | 30,8 | 5,8 | 17 | 1 | IV | 0 | 1 | 5,11,13,21,72 |  |
| *Platyrrhinus lineatus* | 69 | 42,5 | 23 | 13 | 0 | FV | 1 | 0 | 5,11,13,21,72,98 |  |
| *Plecotus auritus* | 42,5 | 39,2 | 8 | 30 | 0 | IV | 0 | 1 | 11,13,21,72 |  |
| *Plecotus austriacus* | 49,5 | 40 | 8 | 26 | 0 | IV | 0 | 1 | 5,11,13,21,<br>72 |  |
| *Plecotus ognevi* | 49 | 41 | 8 | 7,5 | 0 | IV | 0 | 1 | 5,11,13,21 |  |
| *Pteropus alecto* | 233 | 172 | 735 | 20 | 0 | FV | 1 | 0 | 11,13,21,72 |  |
| *Pteropus dasymallus* | 208 | 136 | 490 | 24 | 0 | OM | 1 | 0 | 13,21,72 |  |
| *Pteropus giganteus* | 234 | 170 | 1100 | 40 | 0 | FV | 1 | 0 | 13,21,72 |  |
| *Pteropus hypomelanus* | 210 | 135 | 360 | 20 | 0 | FV | 1 | 0 | 5, 11,13,21,<br>72,99 |  |
| *Pteropus livingstonii* | 340 | 167 | 588 | 15 | 0 | FV | 1 | 0 | 11,13,21,72 |  |
| *Pteropus lylei* | 200 | 153 | 435 | 20 | 0 | FV | 1 | 0 | 5,11,13,21,<br>72 |  |
| *Pteropus niger* | 230 | 156 | 460 | 20 | 0 | FV | 1 | 0 | 5,11,13,21 |  |
| *Pteropus poliocephalus* | 250 | 164 | 855 | 24 | 0 | FV | 1 | 0 | 5,11,13,21,<br>72 |  |
| *Pteropus pumilus* | 168 | 104 | 173 | 17 | 0 | FV | 1 | 0 | 5,11,13,21,<br>72 |  |
| *Pteropus rodricensis* | 175 | 125 | 278 | 28 | 0 | OM | 1 | 0 | 5,11,13,21,<br>72 |  |
| *Pteropus scapulatus* | 161 | 130 | 429 | 16 | 0 | FR | 1 | 0 | 13,21,72 |  |
| *Pteropus vampyrus* | 258 | 203 | 1050 | 24 | 0 | FV | 1 | 0 | 13,21 |  |
| *Rhinolophus euryale* | 63,5 | 55 | 20,8 | 13 | 0 | IV | 0 | 1 | 13, 21,<br>23,72 |  |
| *Rhinolophus ferrumequinum* | 62,5 | 56 | 28,5 | 31 | 0 | IV | 0 | 1 | 13, 21,<br>23,72 |  |
| *Rhinolophus hipposideros* | 41,8 | 38,8 | 6,8 | 29 | 0 | IV | 0 | 1 | 13, 21, 23, 72 |  |
| *Rhinolophus megaphyllus* | 50 | 48 | 9,3 | 7,5 | 0 | IV | 1 | 1 | 5, 11, 13, 21, 72 |  |
| *Roussettus aegyptiacus* | 165 | 94 | 126 | 25 | 0 | FV | 1 | 0 | 5, 11, 13, 21, 100 |  |
| *Roussettus leschenaultii* | 103 | 86,7 | 89,5 | 14 | 0 | FV | 1 | 0 | 5, 11, 13, 21, 72 |  |
| *Saccopteryx bilineata* | 51,5 | 46 | 7,65 | 6 | 0 | IV | 1 | 0 | 13, 21, 23, 72, 101 |  |
| *Scotorepens balstoni* | 51 | 36,3 | 10,2 | 9,7 | 1 | IV | 1 | 0 | 13, 21 |  |
| *Sturnira lilium* | 62 | 42,5 | 20 | 12 | 0 | FV | 1 | 0 | 5, 11, 13, 21, 72 |  |
| *Tadarida brasiliensis* | 54 | 41,5 | 11,5 | 12 | 0 | IV | 0 | 0 | 13, 21, 23, 72 |  |
| *Tadarida teniotis* | 84,5 | 60 | 30 | 13 | 0 | IV | 0 | 0 | 13, 21, 23, 72 |  |
| *Taphozous georgianus* | 70,8 | 70 | 40,5 | 4 | 0 | IV | 1 | 1 | 13, 21 |  |
| *Taphozous hilli* | 72,1 | 67,5 | 22,5 | 8,8 | 0 | IV | 1 | 1 | 13, 21 |  |
| *Vespertilio murinus* | 51 | 45,6 | 17 | 15 | 1 | IV | 0 | 1 | 13, 21, 23, 72, 102 |  |

